# Chronic Chemogenetic Manipulation of Ventral Pallidum Targeted Neurons in Rats Fed an Obesogenic Diet

**DOI:** 10.1101/2020.02.18.954677

**Authors:** Wilder T. Doucette, Elizabeth B. Smedley, Metztli Ruiz-Jaquez, Jibran Y. Khokhar, Kyle Smith

**Author notes:** Corresponding Author: Wilder T. Doucette, MD, PhD, Department of Psychiatry, 1 Medical Center Drive, Lebanon, NH 03756.

## Abstract

Current treatments for obesity are unable to reliably reduce weight over time and alternative treatments need to address this challenge. New interventions that target the nervous system could manipulate brain networks underlying reward, metabolic rate and behavior. Here, the ventral pallidum was evaluated as a target of manipulation due to its established role in the networks underlying motivation, pleasure and behavioral output. Chronic inhibitory or excitatory chemogenetic activation was used to modulate the activity of ventral pallidum (VP) targeted neurons in rats on an obesogenic diet. We hypothesized that inhibition of VP activity would decrease the salience of the rats’ high-sugar, high-fat diet and lead to reduced food consumption and weight gain over time. Paradoxically, measurements of weight, water and food consumption revealed significantly increased weight gain in both groups receiving VP targeted manipulation that was not readily explained by food or water consumption. We theorize that the complex reciprocal feedback between ventral striatal structures (e.g., VP) and metabolic centers of the hypothalamus and brainstem, demonstrated by prior research, likely underpin our findings. This study suggests that the treatment of appetitive disorders (e.g., obesity) with chronic neuromodulation-based interventions could be burdened by the delayed onset of outcomes that are difficult to predict from related prior studies that used acute interventions.

## Introduction

The prevalence of overweight and obese (BMI ≥ 30 kg/m^2^) individuals is increasing at a startling rate despite current treatment efforts, and is considered to be a global epidemic by the World Health Organization [1]. Up to 57.8% of the global adult population is predicted to be overweight or obese by 2030 [2]. Current treatments (e.g., lifestyle interventions, pharmacotherapy and surgical interventions) all suffer from the achieved weight loss not being sustained over the long term in many patients [3-5]. Because of the established morbidity associated with obesity (e.g., metabolic, endocrine and cardiovascular diseases) [6] there is a pressing need to find new and effective treatments.

Basal metabolic rate and behavioral actions (e.g., diet and exercise) are key factors in obesity and result from brain-based neural processes. Thus, interventions to directly manipulate these underlying neural processes are being actively pursued as potential therapeutic strategies. Networks involving reward and motivational processes (e.g., networks involving ventral striatal regions), and regions underpinning energy homeostatic processes (e.g., the hypothalamus and brainstem), have received particular attention. Manipulations of these areas to curb weight gain have included lesions, pharmacologic modulation, and molecular or electrical stimulation [7-9]. Unfortunately, the manipulation of hypothalamic activity to suppress appetite and/or increase energy expenditure have, like other clinical interventions, proven to be rather inadequate over the long term [10, 11]. It has been hypothesized that modulating the mesolimbic reward network could overcome this limitation, and studies using network-based interventions (deep brain stimulation) in preclinical models and patients with obesity have demonstrated potential but with similar findings of time-limited effectiveness and variable effectiveness across individuals [10, 12]. In short, long-term treatment options based on brain intervention remain elusive.

The newest generation of neuroscience technologies has allowed access to neural signaling mechanisms at very fine levels of detail, and many have shown promise in the realm of overeating and weight control [13, 14]. Among these, perhaps the most promising is a chemogenetic approach using Designer Receptors Exclusively Activated by Designer Drugs (DREADDs) [15]. DREADDs offer a naturalistic means of changing neuronal activity through endogenous intracellular signaling mechanisms and a relative non-invasiveness of delivering receptor ligands systemically once brain expression of the DREADD receptor has occurred [16-18]. As a result it has emerged as a highly attractive method for chronic brain manipulation with translational potential.

In most prior studies, acute chemogenetic manipulation produces a limited time window of effect (hours) and is tested on a behavior, such as an eating bout, that can be assessed acutely [16-18]. In contrast, treatment of chronic brain-based disorders (e.g., obesity and psychiatric illnesses) will likely require ongoing modulation on the order of months to years. However, only recently has research started to investigate the effectiveness of chronic chemogenetic manipulation of brain activity with longitudinal behavioral assessment [19,20]. Here, we report an initial attempt to assess the effects of chronic inhibition or activation of ventral pallidum (VP) targeted neurons.

Specifically, we evaluated the effect of chronic inhibitory or excitatory modulation of VP targeted neurons in rats on an obesogenic diet. The VP was selected as a target due to its critical role within the mesolimbic network in regulating motivation, effort, and hedonic processes [21-23]. Notably, drastic damage to VP activity can completely abolish how wanted and liked a palatable food is, the only such brain area to carry this function to our knowledge [21,24]. More subtle and acute inhibitions of the VP can reduce palatable food consumption, while other VP modulations such as those that stimulate the VP can elevate food consumption [21-23,25-28]. Moreover, DREADD-mediated inhibition of the VP can be effective at reducing motivational attraction to food cues [29] and motivation to seek other rewards [25,30,31]. Based on this background, we used an implantable osmotic minipump to deliver clozapine-N-oxide (CNO), a ligand for the DREADD receptor, continuously over ∼1 month and monitored food and water consumption along with weight in rats with expression of hM4Di (Gi - inhibitory DREADD) or hM3D (Gq - excitatory DREADD) as well as a control virus targeted to the VP. We hypothesized that chronic activation of inhibitory Gi receptors targeted to the VP would decrease food consumption and weight compared to the control group, while chronic excitatory Gq receptor activation would increase weight and food consumption.

## Methods and Materials

Male Sprague-Dawley rats (N=30) were purchased from Charles River (Shrewsbury, MA) at postnatal day (PND) 28 and individually housed using a 12 hour light/dark schedule with a palatable high-fat, high-sugar diet (“sweet-fat diet”), which contained 19% protein, 36.2% carbohydrates, and 44.8% fat by calories and 4.6 kcal/g (Teklad Diets 06415, South Easton, MA) and water available *ad libitum*. We used an early exposure to a palatable high calorie diet (starting after weaning from maternal milk) to mirror the US obesity epidemic, in which high-calorie food consumption can emerge in childhood and can lead to a lifetime of treatment resistant obesity [32-35]. Two cohorts of animals were run separately and the data was combined for final analysis. Food, water and weight (all in grams) were manually monitored daily. All experiments were carried out in accordance with the NIH Guide for the Care and Use of Laboratory Animals (NIH Publications No. 80-23) revised in 1996, and were approved by the Institutional Animal Care and Use Committee at Dartmouth College.

### DREADD virus injection

Three groups of animals (N=10 each; PND 70) received bilateral VP viral vector infusions under anesthesia with isoflurane gas using a stereotaxic apparatus (Stoelting, Kiel, WI, USA). Surgery was conducted under aseptic conditions. A 5 µl, 33-gauge beveled needle-tipped syringe (World Precision Instruments, Sarasota, FL, USA) was lowered to the bilateral target sites, in mm: VP −0.12 AP, ± 2.4 ML, −8.2 DV [36] and allowed to rest for 3 minutes prior to infusion. Viral vectors were infused at a rate of 0.15 µl/min. Dispersion of virus was allowed for 5 minutes post-infusion. The following vectors were used, each given at a total volume of 0.8 μl per hemisphere: 1) inhibitory Gi DREADD (AAV5-hSyn-hM4Di-mCherry; UNC Vector Core; n = 5; AAV8-hSyn-hM4Di-mCherry; Addgene; n = 5); excitatory Gq DREADD (AAV8-hSyn-hM3Gq-mCherry; Addgene; n = 10); and control construct (AAV5-hSyn-EYFP; UNC Vector Core; n = 10). Alzet osmotic pumps (model 2ML4; Durect Corp., Cupertino, CA), were implanted subcutaneously (peri-scapular) at PND 90 to deliver CNO (∼6 mg/kg/24 hours, dissolved in saline at PH 4) in all groups until removal at PND 120 (Fig. 1A). Animals received ketoprofen (3 mg/kg) following virus injection and pump implant surgery. Pumps were evaluated post-explant. We confirmed in each case that the CNO solution was evacuated from the pumps and that CNO had not precipitated out of solution within the pump.

**Figure 1.**
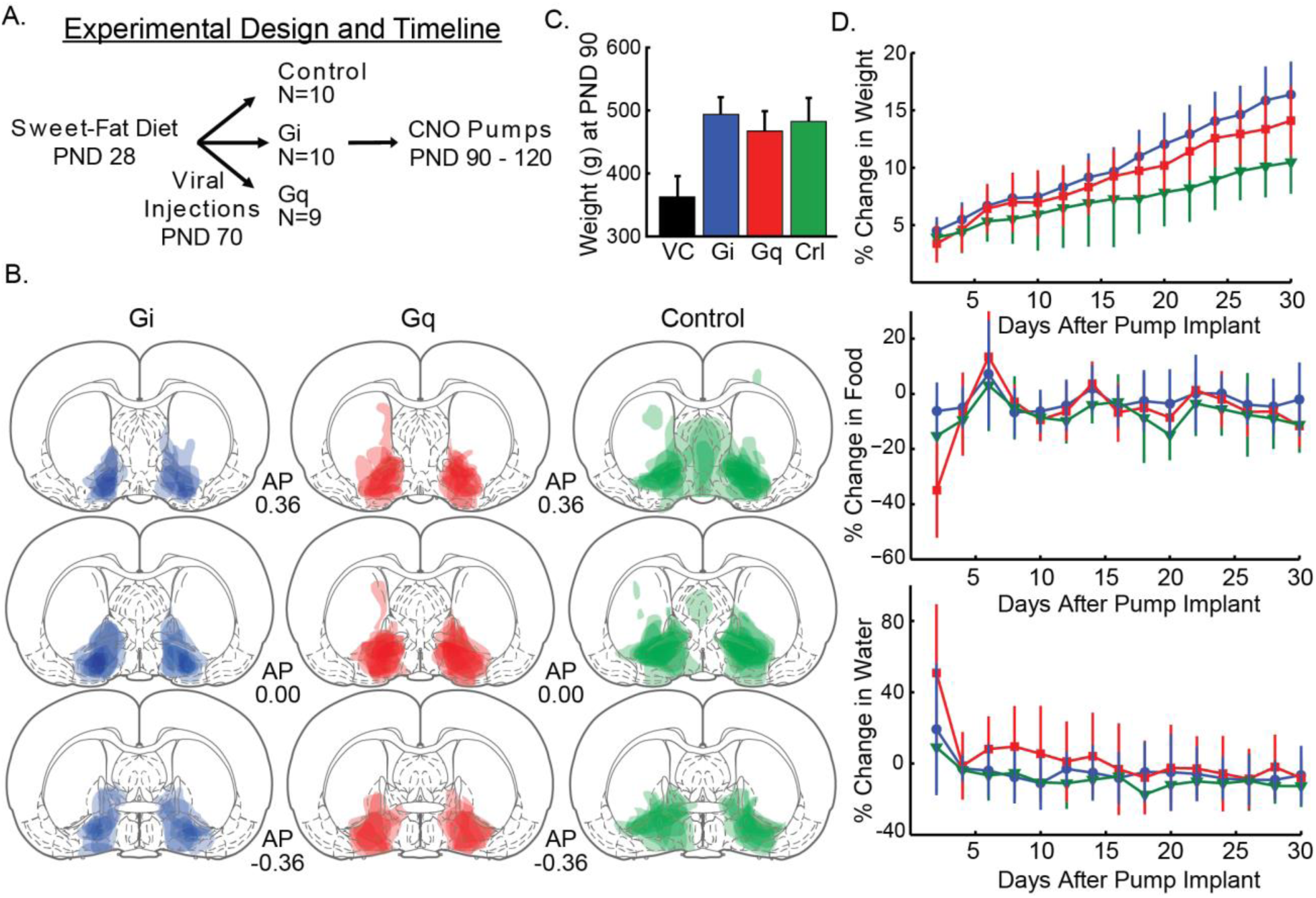
Chronic activation of Gi and Gq in ventral pallidum targeted neurons increases weight gain. Panel **A** illustrates the experimental design and timeline. **B**. The expression of the fluorescent marker mCherry/eYFP for each animal is overlaid in three representative sections along the A-P axis (Bregma: 0.36, 0.00, −0.36) with a separate column and color for each group (Gi - Blue, Gq - Red, control - Green). **C.** Average weight (grams) of the three cohorts (Gi, Gq, and control [crl]) at postnatal day (PND) 90 ± 1 standard deviation. For comparison the average weight of rats on normal chow at PND 90 from the vendor (VC), Charles River, is shown in black. **D.** The effect of chronic CNO delivery on weight, food and water consumption is displayed with line color matching that of panel **B**. Days are graphed as two-day averages. Error bars = ±1 standard deviation.

### Histology

After completion of the weight and intake measurement phase, rats were deeply anesthetized with 1-2 ml of Euthasol (phenobarbital) and perfused with 0.9% saline solution for approximately 6-8 min followed by 10% formalin in distilled H20 until fixture of head and neck tissue (approximately 3-4 min). Brain tissue was extracted, saturated with 20% sucrose (in distilled water) and frozen to −80°C. Brains were then sliced to 60 µm thick sections with a microtome, mounted to slides, and cover-slipped with Vectashield mounting medium containing DAPI. Fluorescent expression was imaged using a microscope (Olympus U-HGLGPS). For each brain, the area of neuronal fluorophore expression was manually transcribed onto blank, printed atlas pages [36] and then transcribed digitally via PowerPoint (Microsoft). Per-animal expression maps were then combined into group expression maps by digitally overlaying the expression areas at 34% transparency (Adobe Illustrator).

### Weight and Intake Measurement and Analyses

Recordings of animal weights began on PND 28 immediately after weaning and continued through to the end of the study. Viral vector infusions occurred on PND 70 and CNO-containing pump implants were placed on PND 90 and per-animal weights, water consumption, and food consumption was measured for the following 30 days until pump removal at PND 120. Measurements were not taken on occasion, but never exceeded two consecutive days (i.e. there are never instances of 3 days passing without data collection). Linear mixed models (below) are well suited for longitudinal studies with missing observations and uneven measurement intervals [37] and thus were chosen for analysis. Analyses include both fixed effects (independent variables; i.e. group, day, and other covariates) as well as random effects (individual rat growth rates and individual starting values; i.e. slopes and y-intercepts). All dependent measures are reported as percent change (%Δ) from a 3-day averaged baseline that was established before pump implantation and chronic CNO administration.

In order to statistically analyze weight, food consumption and water consumption, a linear mixed model was used. The Group variable had three levels (Control, Gi, and Gq), was dummy coded, and compared each experimental group to the mutual control, such that the Gi Group was compared to the Control Group and the Gq Group was compared to the Control Group. All linear mixed models were fit by maximum likelihood and t-tests used Satterthwaite approximations of degrees of freedom (R; “lmerTest”) [38]. The reported statistics include parameter estimates (β values) in the units of the dependent variable, confidence intervals (CI; 95% bootstrapped confidence intervals around dependent variable), standard error of the parameter estimate (SE), and p-values. All statistical tests were carried out using R [39]. Graphs were created with Matlab (R2017b).

## Results

### Viral expression

All animals were evaluated for viral expression in the VP. One animal with expression outside the targeted area was excluded from analysis. Thus, analysis was run with group sizes as: 10 animals in Gi Group, 9 animals in Gq Group, and 10 animals in the Control Group. VP expression was defined by the Paxinos and Watson (2006) atlas, with the target coordinates being ventral to anterior commissure and lateral to the substantia innominata and lateral preoptic area but dorsal to magnocellular preoptic nucleus (coordinates above). The majority of

Gi Group expression was most dense within our area of interest in the VP similar to what we have observed in previously published studies [29,40]. There was some minor/inconsistent spread anterior and medial to the target area (Fig. 1B, left). The Gq Group showed dense and mostly restricted expression in the VP, with some minor dorsal spreading (Fig. 1B, center). The Control Group exhibited more spread outside of the VP target (Fig. 1B, right).

### Percent change in weight from baseline

Fig. 1C illustrates that all three groups with *ad libitum* access to the sweet-fat diet had average weights that were both consistent with prior models of obesity at PND 90 [41], and, according to vendor data [42], likely weighed more than age matched rats from the vendor colony that were fed a standard rat chow. Both the Gi Group and the Gq Group differed in weight from the Control Group by PND 120 (pump removal). The effects on weight did not emerge until over a week into treatment, suggesting that the relevant changes in brain activity (in VP and wider circuitry) occurred over a long time course. Thus, chronic activation must have altered the network of networks connected to the VP in a different way then what was happening in the first 24-48 hours. In general, animals that ate more tended to weigh more, but water consumption levels did not vary with weight gain. <u>While all animals gained weight over the course of the 30 days, the Gi Group and Gq Group gained weight at a significantly faster rate than the control group.</u> **All groups received CNO.**

To reveal this, a linear mixed model was constructed using individual rat percent change in weight from baseline (pre-pump) (%Δ weight; Fig. 1D, top) by fixed effects of Group assignment (Gi, Gq, Control), day (1-30), percent change in food consumption from baseline, and percent change in water consumption from baseline and random effects of slope (i.e. growth curves) and intercept (i.e. individual y-intercepts). There was a significant Group Gi by day interaction (est: 0.161 %Δ weight; CI: 0.06-0.25; SE: 0.049; p = 0.003). A similar difference was found for Group Gq, as the interaction between Group Gq and Group Control by day was significant (est: 0.112 %Δ weight; CI: 0.00-0.21; SE: 0.050; p = 0.033).

There was a significant main effect of day (est: 0.254 %Δ weight; CI: 0.19-0.33; SE: 0.035; p < 0.001) indicating that all animals displayed weight increases by the end of the study. A significant main effect of food consumption (est: 0.014 %Δ weight; CI: 0.01-0.02; SE: 0.003; p < 0.001) shows that animals who weighed more tended to eat more. However, percent change in weight did not vary in accordance with percent change in water consumption (est: −0.002 %Δ weight; CI: −0.01-0.00; SE: 0.003; p = 0.585), suggesting that there were not weight-associated differences in water drinking. The percent change in weight from baseline took time to diverge between the Gi and Gq Groups and the Control Group. Therefore, it was not surprising that neither the Gi Group (est: −0.216 %Δ weight; CI: −1.94-1.64; SE: 0.967; p = 0.825) nor the Gq group (est: −0.184 %Δ weight; CI: −1.84-1.87; SE: 0.993; p = 0.854) differed from the Control Group in percent change in weight measures (on average) across the 30 days.

### Percent change in food consumption

Even though the Gi and Gq Groups gained weight faster, the Groups did not differ in their food consumption compared to controls. Moreover, all animals lessened their overall daily consumption by the end of the study. Animals who ate more tended to drink more and weigh slightly more, though there was no effect of VP perturbation on this relationship. Thus, the elevated weight gain in Gi- and Gq-treated animals was not related to an elevation in the amount of food that they ate. The rats in the Gi Group did not display differences from rats in the Control Group in the amount of food consumed relative to baseline amounts over time as indicated by a non-significant group by day interaction (est: −0.102 %Δ food; CI: −0.46-0.24; SE: 0.174; p = 0.905). The Gq Group also did not differ in consumption from Controls over time as seen in a non-significant group by day interaction (est: −0.175 %Δ food; CI: −0.58-0.17; SE: 0.173; p = 0.429). Further, on average across the 30 days, neither the Gi Group (est: 2.032 %Δ food; CI: −6.41-10.5; SE: 4.363; p = 0.645) nor the Gq Group (est: 2.185 %Δ food; CI: −7.05-11.6; SE: 4.450; p = 0.627) differed from the Control Group in the relative amount of food consumed compared to baseline.

A significant main effect of day (est: −0.456 %Δ food; CI: −0.75-(−0.15); SE: 0.142; p = 0.002) showed that all animals exhibited a relative decrease in food consumption compared to baseline by the end of the study. There was a significant main effect of weight (est: 1.248 %Δ food; CI: 0.64-1.86; SE: 0.302; p < 0.001), indicating that on days when animals had a larger percent change in weight they also had corresponding relative changes in food consumption independent of Group. This correlation was also observed with water consumption (est: 0.125 %Δ food; CI: 0.06-0.19; SE: 0.031; p < 0.001), indicating that from day to day rats had changes in water consumption that related to changes in food consumption independent of Group.

### Percent change in water consumption

Rats from all three Groups tended to drink less over time and water consumption did not fluctuate with weight gain. Again, however, water consumption and food consumption were related as animals tended to drink more if they ate more. The interaction between the Gi Group and Control Group by day was not significant (est: 0.126 %Δ water; CI: −0.27-0.56; SE: 0.236; p = 0.596). In addition, on average across the 30 days, the Gi Group did not significantly differ from the Control Group in relative water consumption levels compared to baseline (est: 0.713 %Δ water; CI: −12.6-12.9; SE: 7.28; p = 0.923). The interaction between the Gq Group and Control Group by day was significant (est: −0.589 %Δ water; CI: −1.04-(−0.14); SE: 0.236; p = 0.018), with Gq animals drinking less over time. However, on average across the 30 days, the Gq Group significantly differed from the Control Group in overall water consumption (est: 19.5 %Δ water; CI: 5.77-33.7; SE: 7.42; p = 0.014). Thus, the Gq Group had increased water consumption relative to baseline compared to the Control Group when averaged across the entire 30 days, despite a lot of variance in the data. However, the Gq Group tended to drink less as time went on compared to Controls in contrast to the time course of relative weight change.

Daily water consumption had decreased relative to baseline amounts by the end of study as seen in a significant main effect of day (est: −0.391 %Δ water; CI: −0.72-(−0.01); SE: 0.192; p = 0.047). Water consumption did not fluctuate with weight gain as suggested by a non-significant main effect of weight (est: −0.012 %Δ water; CI: −0.83-0.79; SE: 0.402; p = 0.977). Food and water consumption were related with a significant main effect of food (est: 0.170 %Δ water; CI: 0.09-0.26; SE: 0.043; p < 0.001).

## Discussion

The goal of this study was to evaluate the therapeutic potential of chronic DREADD activation as a therapy for a chronic condition, obesity. Our results indicate all animals gain weight over time when provided with a high-fat, high-sugar diet consistent with previous findings [34], but animals who received chronic activation of an inhibitory or excitatory DREADD receptor targeted to the VP gain weight at a faster rate than the Control Group. This result is not readily explained by food or water consumption differences between the experimental groups and the control group. The observed differences in weight trajectories that emerged without changes in water or caloric intake has been previously reported when chronic deep brain stimulation was targeted to either the nucleus accumbens [43] or the lateral hypothalamus [44] in models of obesity.

It was surprising that putatively inhibiting VP activity (with Gi DREADDs) did not cause reductions in food intake and/or weight because acute inhibition of the VP are effective at reducing rewarding food consumption, hedonic reactions to food, and motivation to procure food [21-23,25]. Unexpectedly, chronic Gq activation did not increase food intake, even though acute VP excitation is closely linked to such changes [21,45]. There were differences between the control group without DREADDs receiving CNO and those with DREADDs receiving CNO. Thus, DREADD expression plus CNO and not CNO alone was responsible for altering weight without altering food intake. While the cited acute manipulations involved behaviors with food rewards, they did not look at ad libitum access or weight over time and so direct comparisons of acute and chronic DREADD manipulation cannot be made. While our data is not ideal for making claims about acute to chronic DREADD manipulations it is surprising that food consumption was not altered in either direction given the prior acute studies. Overall this highlights the need for careful pre-clinical exploration of chronic brain manipulation where the premise may be based on acute interventions.

Our attempted translational intervention using chronic manipulation of the VP used the same DREADD receptors and CNO ligand that were used in previous acute VP DREADD manipulations that altered VP activity, changed food seeking and eating behavior [16-18]. It is possible that chronic activation caused unexpected changes in DREADD receptor function, CNO (or its clozapine metabolite) efficacy at the DREADD receptors over long periods of administration, CNO metabolism, and/or receptor/intracellular signaling adaptations to chronic agonist activity [46,47]. However, regardless of how the activity of VP targeted neurons was altered, the net result was an increase in weight with no change in caloric intake.

Our surprising outcome could be the result of known reciprocal connectivity between regions within the ventral striatum (VP and nucleus accumbens) and the hypothalamus. The VP has both direct connections with the lateral hypothalamus [48], and indirect connections by way of the nucleus accumbens shell [49-51] that also has connection with the lateral hypothalamus [52,53]. Given the strong connectivity between the ventral striatal structures and the hypothalamus it is not surprising that chronic manipulation (deep brain stimulation - DBS) of either regions within the ventral striatum or the hypothalamus can produce similar effects on metabolism or motivated behavior. DBS of the nucleus accumbens shell or the lateral hypothalamus have both produced weight changes in rodent models of obesity without behavioral effects on food or water intake [43,44]. Moreover, Diepenbroek et al. showed that DBS of the nucleus accumbens shell could produce changes in peripheral glucose metabolism [54]. Both human ventral striatal DBS in a patient with obsessive compulsive disorder and optogenetic activation of dopamine D1 receptor neurons in nucleus accumbens of mice have been shown to increase glucose tolerance and insulin sensitivity [55]. Manipulation of lateral hypothalamic neuronal populations are known to alter both basal metabolic rate and spontaneous physical activity [56] and could be possible mediators of our DREADD manipulation by altering energy homeostatic processes that produced the observed weight changes.

Several features of the study design limit the breadth of conclusions that can be drawn from the presented results. There were no cohorts fed a “control” rat chow and prior work has shown that rats with ad libitum access to standard chow also develop obesity [57]. Future evaluation of a lower caloric, less palatable food would provide insight into the influence of food salience on the effect of chronic VP manipulation using DREADDs but would have less relevance to the treatment of obesity. Even if the primary outcome (weight reduction) was achieved, resolving what the relevant activity changes were (within the DREADD expression areas and elsewhere in the brain), while important, would be a truly substantial undertaking and one that might ultimately not be realistic to fully do. Those changes might include neuronal firing activity changes, DREADD receptor function and distribution changes, CNO binding or metabolism changes, intracellular signaling changes, etc. While viral injections were targeted to the VP and this was the structure with the highest density of expression, there was expression in surrounding brain regions that could also have contributed to the study outcome. It is important to note that the degree of off target expression within our Gi and Gq cohorts was similar to those observed in prior published studies from our group that have used acute DREADD manipulations [29,40]. It is possible that more localized, cell type specific, or connectivity specific expression of DREADDS within the VP could produce translationally relevant changes in food consumption and weight [51].

## Conclusion

Our results are the first to show that attempts to chronically modulate VP activity in rats on an obesogenic diet do not alter food consumption as hypothesized from studies that have used acute chemogenetic manipulations. Instead, chronic bidirectional VP manipulation paradoxically increased weight gain without changing food or water consumption. Our findings highlight the challenges of translating the findings from acute chemogenetic studies into treatments for chronic neuropsychiatric illnesses.

## Acknowledgements

The authors would like to acknowledge Alyssa C. DiLeo and Alex R. Brown for their help with histology.

## References

1. James WP (2008) WHO recognition of the global obesity epidemic. Int J Obes (Lond) 32 Suppl 7: S120–126.

2. Gonzalez-Muniesa P, Martinez-Gonzalez MA, Hu FB, Despres JP, Matsuzawa Y, et al. (2017) Obesity. Nat Rev Dis Primers 3: 17034.

3. Christou NV, Look D, Maclean LD (2006) Weight gain after short- and long-limb gastric bypass in patients followed for longer than 10 years. Ann Surg 244: 734–740.

4. Khera R, Murad MH, Chandar AK, Dulai PS, Wang Z, et al. (2016) Association of Pharmacological Treatments for Obesity With Weight Loss and Adverse Events: A Systematic Review and Meta-analysis. JAMA 315: 2424–2434.

5. Gotthardt JD, Bello NT (2016) Can we win the war on obesity with pharmacotherapy? Expert Rev Clin Pharmacol: 1–9.

6. Heymsfield SB, Wadden TA (2017) Mechanisms, Pathophysiology, and Management of Obesity. N Engl J Med 376: 254–266.

7. Teitelbaum P, Epstein AN (1962) The lateral hypothalamic syndrome: recovery of feeding and drinking after lateral hypothalamic lesions. Psychol Rev 69: 74–90.

8. Anand BK, Brobeck JR (1951) Localization of a feeding center in the hypothalamus of the rat. Proc Soc Exp Biol Med 77: 323–324.

9. Nangunoori RK, Tomycz ND, Oh MY, Whiting DM (2016) Deep Brain Stimulation for Obesity: From a Theoretical Framework to Practical Application. Neural Plast 2016: 7971460.

10. Whiting DM, Tomycz ND, Bailes J, de Jonge L, Lecoultre V, et al. (2013) Lateral hypothalamic area deep brain stimulation for refractory obesity: a pilot study with preliminary data on safety, body weight, and energy metabolism. J Neurosurg 119: 56–63.

11. Hamani C, McAndrews MP, Cohn M, Oh M, Zumsteg D, et al. (2008) Memory enhancement induced by hypothalamic/fornix deep brain stimulation. Ann Neurol 63: 119–123.

12. Harat M, Rudas M, Zielinski P, Birska J, Sokal P (2016) Nucleus accumbens stimulation in pathological obesity. Neurol Neurochir Pol 50: 207–210.

13. Krashes MJ, Kravitz AV (2014) Optogenetic and chemogenetic insights into the food addiction hypothesis. Front Behav Neurosci 8: 57.

14. Palmiter RD (2017) Neural Circuits That Suppress Appetite: Targets for Treating Obesity? Obesity (Silver Spring) 25: 1299–1301.

15. Armbruster BN, Li X, Pausch MH, Herlitze S, Roth BL (2007) Evolving the lock to fit the key to create a family of G protein-coupled receptors potently activated by an inert ligand. Proc Natl Acad Sci U S A 104: 5163–5168.

16. Smith KS, Bucci DJ, Luikart BW, Mahler SV (2016) DREADDS: Use and application in behavioral neuroscience. Behav Neurosci 130: 137–155.

17. Roth BL (2016) DREADDs for Neuroscientists. Neuron 89: 683–694.

18. Burnett CJ, Krashes MJ (2016) Resolving Behavioral Output via Chemogenetic Designer Receptors Exclusively Activated by Designer Drugs. J Neurosci 36: 9268–9282.

19. Iyer SM, Vesuna S, Ramakrishnan C, Huynh K, Young S, et al. (2016) Optogenetic and chemogenetic strategies for sustained inhibition of pain. Sci Rep 6: 30570.

20. Yu S, Munzberg H (2018) Testing Effects of Chronic Chemogenetic Neuronal Stimulation on Energy Balance by Indirect Calorimetry. Bio Protoc 8.

21. Smith KS, Tindell AJ, Aldridge JW, Berridge KC (2009) Ventral pallidum roles in reward and motivation. Behav Brain Res 196: 155–167.

22. Castro DC, Cole SL, Berridge KC (2015) Lateral hypothalamus, nucleus accumbens, and ventral pallidum roles in eating and hunger: interactions between homeostatic and reward circuitry. Front Syst Neurosci 9: 90.

23. Root DH, Melendez RI, Zaborszky L, Napier TC (2015) The ventral pallidum: Subregion-specific functional anatomy and roles in motivated behaviors. Prog Neurobiol 130: 29–70.

24. Cromwell HC, Berridge KC (1993) Where does damage lead to enhanced food aversion: the ventral pallidum/substantia innominata or lateral hypothalamus? Brain Res 624: 1–10.

25. Chang SE, Smedley EB, Stansfield KJ, Stott JJ, Smith KS (2017) Optogenetic Inhibition of Ventral Pallidum Neurons Impairs Context-Driven Salt Seeking. J Neurosci 37: 5670–5680.

26. Tooley J, Marconi L, Alipio JB, Matikainen-Ankney B, Georgiou P, et al. (2018) Glutamatergic Ventral Pallidal Neurons Modulate Activity of the Habenula-Tegmental Circuitry and Constrain Reward Seeking. Biol Psychiatry 83: 1012–1023.

27. Richard JM, Stout N, Acs D, Janak PH (2018) Ventral pallidal encoding of reward-seeking behavior depends on the underlying associative structure. Elife 7.

28. Creed M, Ntamati NR, Chandra R, Lobo MK, Luscher C (2016) Convergence of Reinforcing and Anhedonic Cocaine Effects in the Ventral Pallidum. Neuron 92: 214–226.

29. Chang SE, Todd TP, Bucci DJ, Smith KS (2015) Chemogenetic manipulation of ventral pallidal neurons impairs acquisition of sign-tracking in rats. Eur J Neurosci 42: 3105–3116.

30. Mahler SV, Vazey EM, Beckley JT, Keistler CR, McGlinchey EM, et al. (2014) Designer receptors show role for ventral pallidum input to ventral tegmental area in cocaine seeking. Nat Neurosci 17: 577–585.

31. Prasad AA, McNally GP (2016) Ventral Pallidum Output Pathways in Context-Induced Reinstatement of Alcohol Seeking. J Neurosci 36: 11716–11726.

32. West DB, Diaz J, Woods SC (1982) Infant gastrostomy and chronic formula infusion as a technique to overfeed and accelerate weight gain of neonatal rats. J Nutr 112: 1339–1343.

33. Morris MJ, Velkoska E, Cole TJ (2005) Central and peripheral contributions to obesity-associated hypertension: impact of early overnourishment. Exp Physiol 90: 697–702.

34. Hariri N, Thibault L (2010) High-fat diet-induced obesity in animal models. Nutr Res Rev 23: 270–299.

35. Cheng HS, Ton SH, Phang SCW, Tan JBL, Abdul Kadir K (2017) Increased susceptibility of post-weaning rats on high-fat diet to metabolic syndrome. J Adv Res 8: 743–752.

36. Paxinos G, Watson C (2006) The rat brain in stereotaxic coordinates. 6th ed. Amsterdam; Boston; Oxford: Academic,. pp. 1 online resource (xxxi p.).

37. Gibbons RD, Hedeker D, DuToit S (2010) Advances in analysis of longitudinal data. Annu Rev Clin Psychol 6: 79–107.

38. Kuznetsova A, Brockhoff B, Christensen P (2016) lmerTest: test in linear mixed effects models. 2.0-33 ed. pp. R package.

39. R-Core-Team (2016) R: a language and environment for statistical computing. R Foundation for Statistical Computing. Vienna, Austria.

40. Chang SE, Todd TP, Smith KS (2018) Paradoxical accentuation of motivation following accumbens-pallidum disconnection. Neurobiol Learn Mem 149: 39–45.

41. Furnes MW, Zhao CM, Stenstrom B, Arum CJ, Tommeras K, et al. (2009) Feeding behavior and body weight development: lessons from rats subjected to gastric bypass surgery or high-fat diet. J Physiol Pharmacol 60 Suppl 7: 25–31.

42. Charles-River-Laboratories (2019). pp. SD Rat Growth Chart.

43. Prinz P, Kobelt P, Scharner S, Goebel-Stengel M, Harnack D, et al. (2017) Deep brain stimulation alters light phase food intake microstructure in rats. J Physiol Pharmacol 68: 345–354.

44. Sani S, Jobe K, Smith A, Kordower JH, Bakay RA (2007) Deep brain stimulation for treatment of obesity in rats. J Neurosurg 107: 809–813.

45. Stratford TR, Kelley AE, Simansky KJ (1999) Blockade of GABAA receptors in the medial ventral pallidum elicits feeding in satiated rats. Brain Res 825: 199–203.

46. Mahler SV, Aston-Jones G (2018) CNO Evil? Considerations for the Use of DREADDs in Behavioral Neuroscience. Neuropsychopharmacology 43: 934–936.

47. Gomez JL, Bonaventura J, Lesniak W, Mathews WB, Sysa-Shah P, et al. (2017) Chemogenetics revealed: DREADD occupancy and activation via converted clozapine. Science 357: 503–507.

48. Faget L, Zell V, Souter E, McPherson A, Ressler R, et al. (2018) Opponent control of behavioral reinforcement by inhibitory and excitatory projections from the ventral pallidum. Nat Commun 9: 849.

49. Zahm DS, Brog JS (1992) On the significance of subterritories in the accumbens part of the rat ventral striatum. Neuroscience 50: 751–767.

50. Zahm DS (2000) An integrative neuroanatomical perspective on some subcortical substrates of adaptive responding with emphasis on the nucleus accumbens. Neurosci Biobehav Rev 24: 85–105.

51. Churchill L, Kalivas PW (1994) A topographically organized gamma-aminobutyric acid projection from the ventral pallidum to the nucleus accumbens in the rat. J Comp Neurol 345: 579–595.

52. Huang XF, Han M, South T, Storlien L (2003) Altered levels of POMC, AgRP and MC4-R mRNA expression in the hypothalamus and other parts of the limbic system of mice prone or resistant to chronic high-energy diet-induced obesity. Brain Res 992: 9–19.

53. Carus-Cadavieco M, Gorbati M, Ye L, Bender F, van der Veldt S, et al. (2017) Gamma oscillations organize top-down signalling to hypothalamus and enable food seeking. Nature 542: 232–236.

54. Diepenbroek C, van der Plasse G, Eggels L, Rijnsburger M, Feenstra MG, et al. (2013) Alterations in blood glucose and plasma glucagon concentrations during deep brain stimulation in the shell region of the nucleus accumbens in rats. Front Neurosci 7: 226.

55. Ter Horst KW, Lammers NM, Trinko R, Opland DM, Figee M, et al. (2018) Striatal dopamine regulates systemic glucose metabolism in humans and mice. Sci Transl Med 10.

56. Zink AN, Bunney PE, Holm AA, Billington CJ, Kotz CM (2018) Neuromodulation of orexin neurons reduces diet-induced adiposity. Int J Obes (Lond) 42: 737–745.

57. Laaksonen KS, Nevalainen TO, Haasio K, Kasanen IH, Nieminen PA, et al. (2013) Food and water intake, growth, and adiposity of Sprague-Dawley rats with diet board for 24 months. Lab Anim 47: 245–256.

